# Interhemispheric Connectivity of the Human Temporal Lobes

**DOI:** 10.1101/2024.07.01.601568

**Authors:** Jeffrey R. Binder, Mónica Giraldo-Chica, Jedediah Mathis, Jia-Qing Tong, Sidney Schoenrock, Volkan Arpinar, Joseph Heffernan, L. Tugan Muftuler

## Abstract

Much is known regarding the major white matter pathways connecting the right and left temporal lobes, which project through the posterior corpus callosum, the anterior commissure, and the dorsal hippocampal commissure. However, details about the spatial location of these tracts are unclear, including their exact course and proximity to cortical and subcortical structures, the spatial relations between corpus callosum and anterior commissure projections, and the caudal extent of transcallosal connections within the splenium. We present an atlas of these tracts derived from high angular resolution diffusion tractography maps, providing improved visualization of the spatial relationships of these tracts. The data show several new details, including branching of the transcallosal pathway into medial and lateral divisions, projections of the transcallosal pathway into the external capsule and claustrum, complex patterns of overlap and interdigitation of the transcallosal and anterior commissure tracts, distinct dorsal and ventral regions of the splenium with high tract densities, and absence of temporal lobe projections in the caudal third of the splenium. Intersection of individual tract probability maps with individual cortical surfaces were used to identify likely regions with relatively higher cortical termination densities. These data should be useful for planning surgical approaches involving the temporal lobe and for developing functional-anatomical models of processes that depend on interhemispheric temporal lobe integration, including speech perception, semantic memory, and social cognition.

**Highlights:**

- Interhemispheric connections of the human temporal lobes were visualized using high angular resolution diffusion tensor imaging tractography.
- Results are displayed on serial orthogonal sections to reveal detailed spatial relationships.
- Corpus callosum projections through the splenium form distinct dorsal and ventral bundles and are absent from the caudal splenium.
- The transcallosal pathway consists of distinct medial and lateral divisions.
- The results reveal projections to the external capsule and claustrum not previously described.
- Transcallosal and anterior commissural pathways show complex patterns of overlap and interdigitation.
- Surface mapping revealed areas with relatively high density of projections to the cortical surface.

## Introduction

Interhemispheric connections of the temporal lobes have been studied extensively in human and nonhuman primate brains. These pathways run through the posterior corpus callosum (CC), anterior commissure (AC), and dorsal hippocampal commissure (DHC) (Demeter, Rosene et al. 1990, Schmahmann and Pandya 2006, Catani and Thiebaut de Schotten 2012). Autoradiographic tracer studies in the macaque monkey suggest broad connectivity throughout much of the temporal lobe, with AC fibers projecting mainly to the rostral third of the temporal cortex, CC fibers mainly to the caudal two-thirds, and DHC fibers to the parahippocampus (Cipolloni and Pandya 1985, Demeter, Rosene et al. 1990, Pandya and Seltzer 1993). Methods used in the human brain have included postmortem staining for axonal degeneration after natural lesions (Di Virgilio, Clarke et al. 1999), blunt dissection of autopsy specimens using the Klingler method (Peltier, Verclytte et al. 2010, Baudo, Colombo et al. 2022), and diffusion imaging tractography (Huang, Zhang et al. 2005, Hofer and Frahm 2006, Park, Kim et al. 2008, Patel, Toussaint et al. 2010, Peltier, Verclytte et al. 2010, Wei, Mao et al. 2017, Archer, Coombes et al. 2019, Baudo, Colombo et al. 2022).

The anatomical relationships of these tracts and their cortical targets in the human brain, however, are still somewhat unclear. Postmortem dissection provides detailed information about the anatomical relations between major tracts and surrounding structures but is limited in its ability to track smaller bundles or discriminate intermingled fibers within a tract. Advances in diffusion imaging have gradually improved the ability to trace smaller tracts, revealing a broader distribution of projections than had been suggested by postmortem dissection (Patel, Toussaint et al. 2010, Peltier, Verclytte et al. 2010, Wei, Mao et al. 2017, Baudo, Colombo et al. 2022). A recent study by Baudo et al. (2022), in particular, provides compelling evidence for projections of the AC to occipital and parietal regions, supporting earlier limited evidence from tractography (Patel, Toussaint et al. 2010) and postmortem axonal degeneration (Di Virgilio, Clarke et al. 1999) studies. Most tractography studies, however, were focused on confirming the general location of major bundles known from postmortem dissection. Results were typically presented as 3D reconstructions that provide only limited information about anatomical relationships and cortical endpoints. Wei et al. (2017) applied advanced diffusion imaging techniques to study CC temporal lobe connections. Their focus was on subdividing regions of the splenium based on lobar projection patterns, and they provided only limited information regarding the location of fibers within the temporal lobe or their terminal projections.

The clearest information on tract location to date comes from a study by Archer et al. (2019), who created a 3D atlas of CC projections called the Transcallosal Tract Template (TCATT) using data from the Human Connectome Project. The atlas includes templates of tracts connecting the CC with superior, middle, and inferior temporal gyri, and a detailed connectivity-based topographic map of the CC. The aim of the TCATT is to provide symmetrical templates for measuring white matter diffusion indices. To achieve this goal, the authors averaged the left and right hemisphere tract maps after reflecting them across the midline, an approach that compromises spatial detail if the left and right tracts are not perfectly symmetric. Archer et al. did not examine temporal lobe connections through the AC, or CC projections to ventral and medial temporal lobe regions.

The goal of the present study was to provide a more complete atlas of interhemispheric pathways within the human temporal lobe white matter, as well as estimates of their principal cortical targets, using high-angular-resolution diffusion imaging (HARDI) tractography. We mapped temporal lobe connections through both CC and AC separately for each hemisphere, and we used a waypoint mask that included the entire temporal lobe. We describe several anatomical features of these pathways that have not been previously noted, and we provide further evidence for the absence of temporal lobe connections in the most caudal aspect of the splenium. The results are presented here as serial sections in orthogonal directions and online as a publicly available digital atlas. This atlas should be useful for planning surgical approaches involving the temporal lobe. It may also contribute to understanding the effects of unilateral temporal lobe lesions on processes thought to engage interhemispheric temporal lobe integration. For example, speech perception impairment after unilateral left temporal lobe damage has been attributed by some authors to a hypothetical disconnection between right hemisphere auditory cortex and left temporal lobe language networks (Takahashi, Kawamura et al. 1992, Poeppel 2001, Maffei, Capasso et al. 2017). Such a mechanism is difficult to assess without detailed information about the white matter location and terminal points of these interhemispheric pathways.

## Methods

### Participants

The participants were 28 healthy, right-handed adults (17 women, 11 men) aged 23 to 50 (mean age 34.6) with no history of neurologic disease. Participants were recruited and scanned at the Medical College of Wisconsin (MCW). All participants provided written informed consent for the study. The project was approved by the MCW Institutional Review Board and was performed in accordance with the Declaration of Helsinki.

### MRI Acquisition

MRI was performed on a 3T GE Premier scanner at the MCW Center for Imaging Research. A 32-channel Nova head receive coil was used for all studies. High-resolution T1-weighted images were acquired using a magnetization prepared gradient echo sequence with 0.8 mm isotropic voxels. High-resolution T2-weighted images were acquired using a 3D Cube sequence, also with 0.8 mm isotropic voxels. Diffusion imaging used spin-echo echoplanar imaging with voxel size = 1.58 x 1.58 x 1.5 mm, TR = 4819 ms, TE = 77.6 ms, multi-band factor = 3, FOV = 221 x 221 mm, matrix = 140 x 140, and 96 axial slices. Data were acquired as 2 shells with b-values of 1000 and 2000 s/mm^2^, each with 75 diffusion sampling directions (listed in Table S1 of Supplemental Information) and either 9 or 10 b0 images, following the HCP Lifespan protocol (Harms, Somerville et al. 2018). Each shell was acquired separately in AP and PA phase-encoding directions to allow distortion correction and subsequent pairwise averaging of each AP-PA image pair. Each of the four runs lasted 6.8 minutes, and total acquisition time was 27.3 minutes.

### DTI Analyses

DWI images were skull-stripped with FSL BET (v 2.1) and de-noised using a spatially adaptive filter (Manjon, Coupe et al. 2010) in ANTs v 2.2.0. Head movement, geometric distortion, and eddy current induced distortions were estimated and corrected using Eddy and Topup tools from FSL (v 6.0.4). Examples of raw and preprocessed images are provided in Figure S1 of Supplemental Information. Average root-mean-square head movement after Eddy correction was 0.384 mm (range 0.20 to 0.97). Outlier detection in Eddy was used to replace (by interpolation) slices exhibiting signal dropout due to movement or other acquisition artifacts. None of the participants had more than 5% outliers (mean ± SD = 2.67% ± 1.17).

Camino (Cook, Bai et al. 2006) was used to generate fractional anisotropy (FA) maps and to estimate SNR on the white matter skeleton of the b0 images as a quality control before running tractography. The SNR values of the sample ranged from 16.6 to 21.0 [mean ± SD = 19.09 ± 1.07]. These values are well above 10, the value below which additional bias from the estimation of the diffusion parameters becomes significant, thus no images were excluded due to low SNR (Pierpaoli and Basser 1996). Additional visual inspection was performed to verify the quality of the data and the correctness of the vectors. Transformation to standard MNI space was done with ANTs v. 2.2.0 and the MNI152_T1_2009c space as target. MNI152_T1_2009c is an average of structural images from 152 healthy adults after high-dimensional nonlinear registration into the MNI152 coordinate space (Fonov, Evans et al. 2009). The transformation from individual FA space was computed by first registering the MNI152_T1_1mm brain template used in FSL to the MNI152_T1_2009c template, which also registers the FMRIB58_FA_1mm FA template to MNI152_T1_2009c space. Individual FA maps were then registered to this version of the FMRIB58_FA_1mm template.

BedpostX was used to estimate diffusion parameters and to model crossing fibers at each voxel. Temporal lobe and corpus callosum ROIs were created for probabilistic tractography of the transcallosal pathways (**Figure 1**). The left and right temporal lobe ROIs included all voxels in the temporal lobe of the MNI152_t1_2009c brain template. The corpus callosum ROI encompassed the entire splenium from the midline to 5 mm left and right of midline (Figure 1b). The inverse transformation obtained from registration to MNI152_T1_2009c space was applied to the three ROIs to run tractography in native space. Two probabilistic tractography maps were generated in each participant using ProbtrackX. One used the right temporal ROI as seed region and the splenium and left temporal ROI as waypoint masks, retaining only the streamlines that passed through both waypoint regions in that order. The second tractography map used the left temporal ROI as seed region and the splenium and right temporal ROI as waypoint masks. This produced two tractography maps containing the number of streamlines reaching each voxel and connecting the three ROIs in the designated order. This method captures both homotopic and heterotopic temporal lobe projections as long as they pass through the corpus callosum ROI. Default stop criteria in ProbtrackX were used for generating streamlines, including stopping at the brain surface OR after traveling 2000 steps (step length = 0.5 mm) OR when streamline curvature exceeded 0.2 (corresponding to the cosine of a minimum angle of approximately ±80 degrees). The probabilities of the two maps when then added to construct the final connectogram of each participant. Connectograms were transformed into standard space using the previously obtained transformation and then averaged across participants. Finally, the average map was thresholded at 0.3% to remove voxels with low mean probability values for easier visualization.

**Figure 1.**
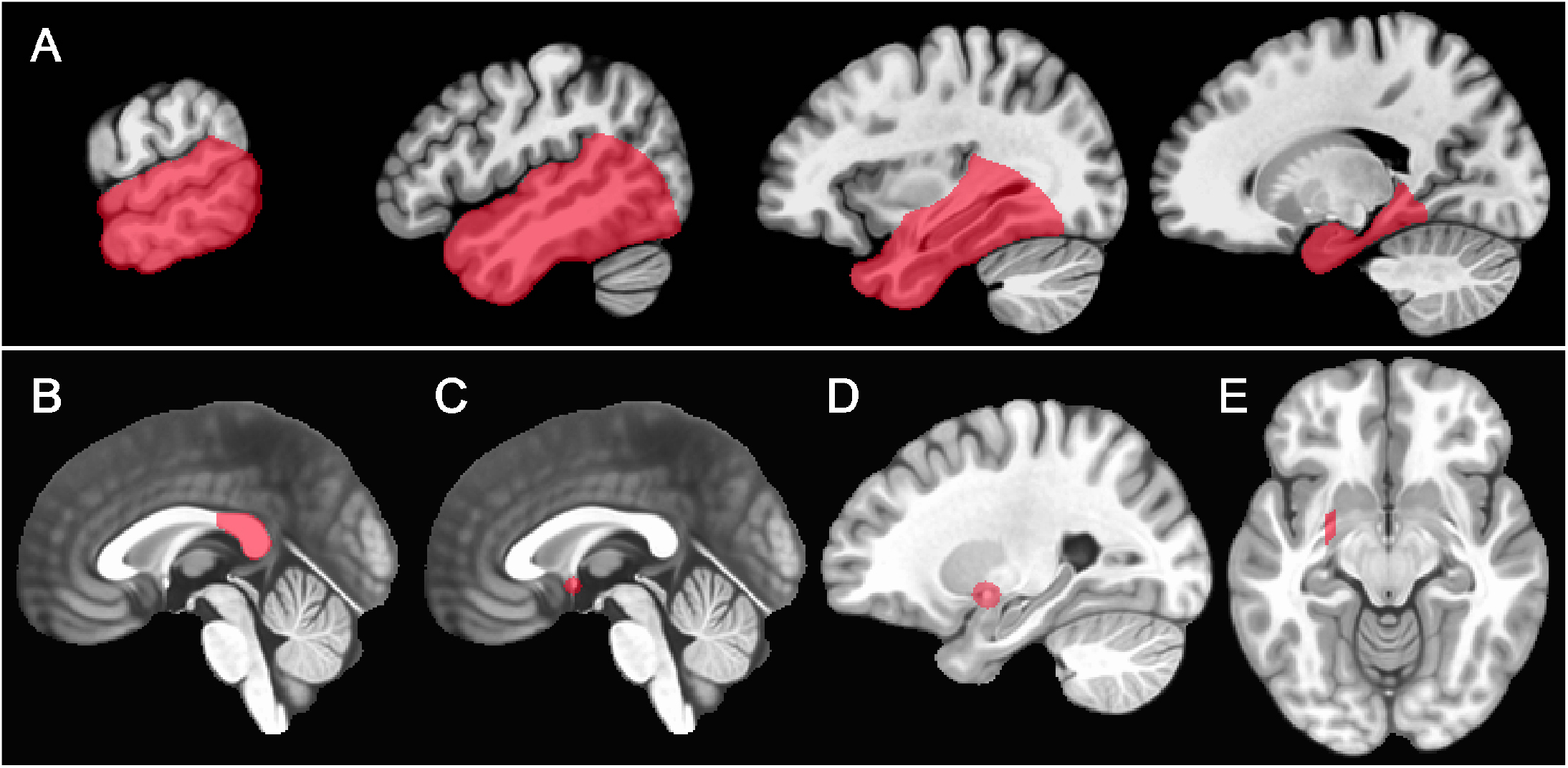
Waypoint regions of interest (ROIs) for tractography. (A) Left temporal lobe ROI used for transcallosal tractography. The right temporal lobe ROI covered the same structures on the right side. (B) Splenium ROI for transcallosal tractography. (C) Anterior commissure ROI. (D, E) Sagittal and axial views of the right Gratiolet canal ROI used for anterior commissure tractography. A symmetric ROI over the left Gratiolet canal was also used.

Temporal lobe connections through the anterior commissure were traced using an 8-mm diameter, 5-mm thick disc-shaped ROI centered on the anterior commissure at the midline (Figure 1c), and two 14-mm diameter, 5-mm thick, vertically oriented, disc-shaped ROIs centered at ±26 mm from the midline to encompass the posterior branch of the anterior commissure as it passes through the Gratiolet canal on each side (Figure 1d-e). A strategy identical to that used for the corpus callosum tracts was followed using these ROIs, producing two tractography maps (from left to right Gratiolet canal via anterior commissure, and vice versa) that were then combined at the individual level and averaged across participants. Using the Gratiolet canal as a waypoint eliminates anterior tract fibers medial to the Gratiolet canal that project to the frontal lobe (Baudo, Colombo et al. 2022).

Finally, average maps of the cortical projections of these interhemispheric connections were created as follows. First, a model of the cortical ribbon of each participant was created from their T1 and T2 images using Freesurfer v7.2. Each participant’s probabilistic tractography map was then intersected with this cortical ribbon prior to any smoothing or thresholding. Individual cortical maps were then registered to a template surface created from the MNI152_t1_2009c brain and smoothed with a 5 mm FWHM kernel constrained to the surface. These unthresholded maps were then averaged across participants. Finally, the values were thresholded at a nominal 0.05% to highlight areas with relatively greater mean probability values.

## Results

### Transcallosal Connections

A detailed map of the temporal lobe transcallosal connections is shown in **Figure 2** on serial sections in three orthogonal planes. As previously described (Huang, Zhang et al. 2005, Hofer and Frahm 2006, Wei, Mao et al. 2017), these tracts cross through the splenium of the corpus callosum. As illustrated in **Figure 3**, they form a somewhat distinct high-probability bundle along the dorsal splenium (midline peak MNI LPI coordinates: 0,-37,20), and a second high-probability bundle in the anterior-inferior splenium (midline peak MNI LPI coordinates: 0,-31,14) that is contiguous with the dorsal hippocampal commissure (DHC). The most caudal sector of the splenium appears to have essentially no temporal lobe connections. Lateral to the midline, the two splenium bundles fuse into a single tract oriented laterally, posteriorly, and slightly superiorly that courses just above the atrium and posterior body of the lateral ventricle. The DHC, in contrast, courses medial to and beneath the atrium to enter the posterior parahippocampus (Gloor, Salanova et al. 1993, Wei, Mao et al. 2017).

**Figure 2.**
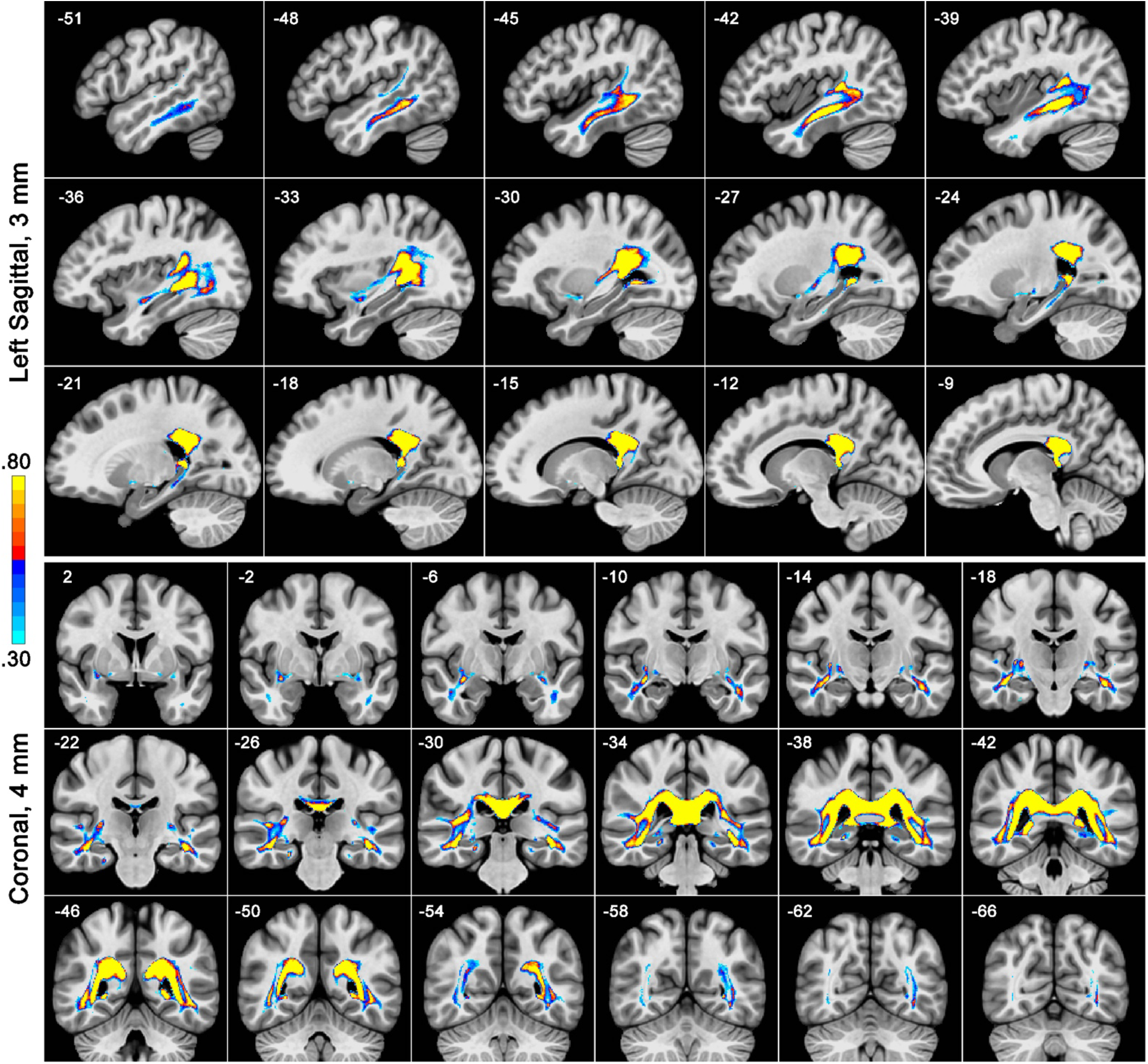
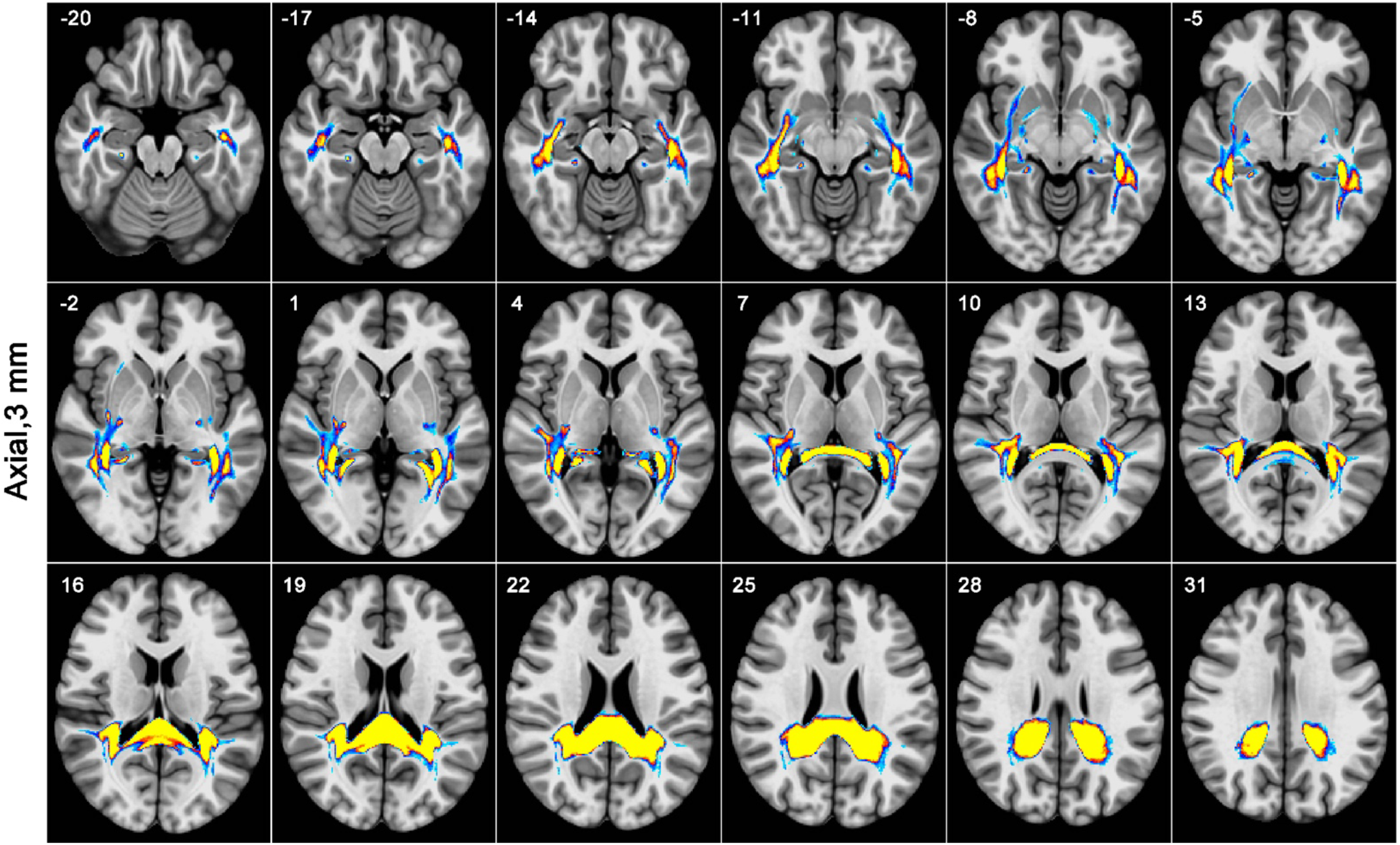
part 1. Serial sections through the averaged group transcallosal tractography map, including left sagittal, coronal, and axial views. Values on the color scale represent % probability. Numbers in each panel indicate the MNI LPI axis location of each slice. The left hemisphere in coronal and axial images is on the reader’s right.

**Figure 3.**
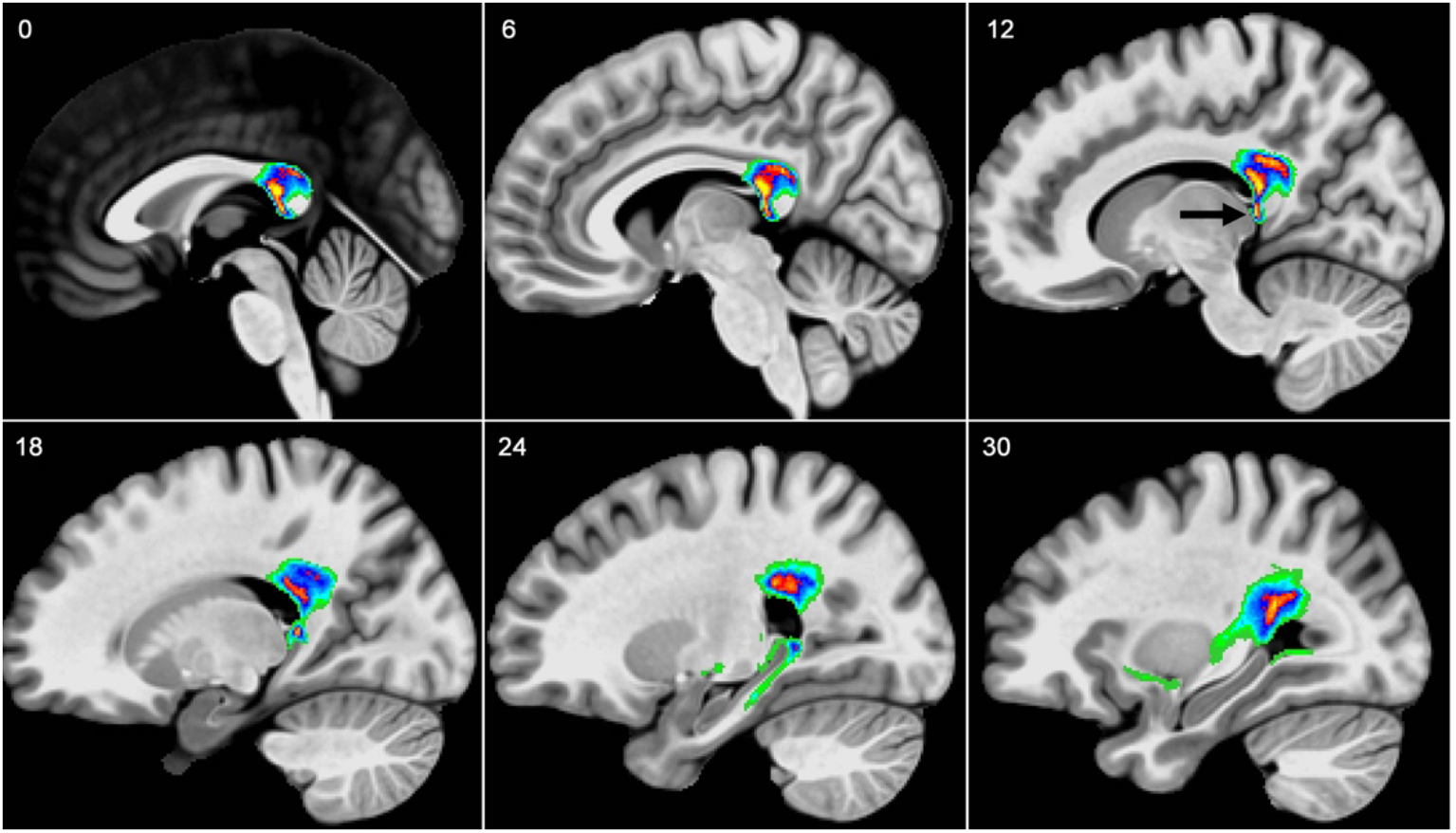
Serial sagittal sections through the right hemisphere transcallosal probability map, highlighting variation in fiber tract probability in the splenium. Distinct dorsal and ventral bundles are especially clear at x = 12 (upper right panel), where a black arrow indicates the dorsal hippocampal commissure (DHC). Lower panels show the DHC coursing into the posterior parahippocampus (x = 18-24) and the dorsal and ventral splenium bundles merging above the atrium.

At the dorsolateral aspect of the atrium, the splenial tract separates into distinct medial and lateral divisions (**Figure 4a**), an arrangement not previously noted. The medial division follows the lateral edge of the atrium within the tapetum, curving anteriorly to enter the sagittal stratum lateral to the temporal horn of the lateral ventricle, where it continues into the deep white matter of the anterior temporal stem, ending at approximately the anterior limit of the stem. Approximately midway along its anterior course it sends a branch, more prominent in the right hemisphere in our sample, that courses superomedially through the stem and enters the inferior aspect of the external capsule and claustrum (**Figure 4b**).

**Figure 4.**
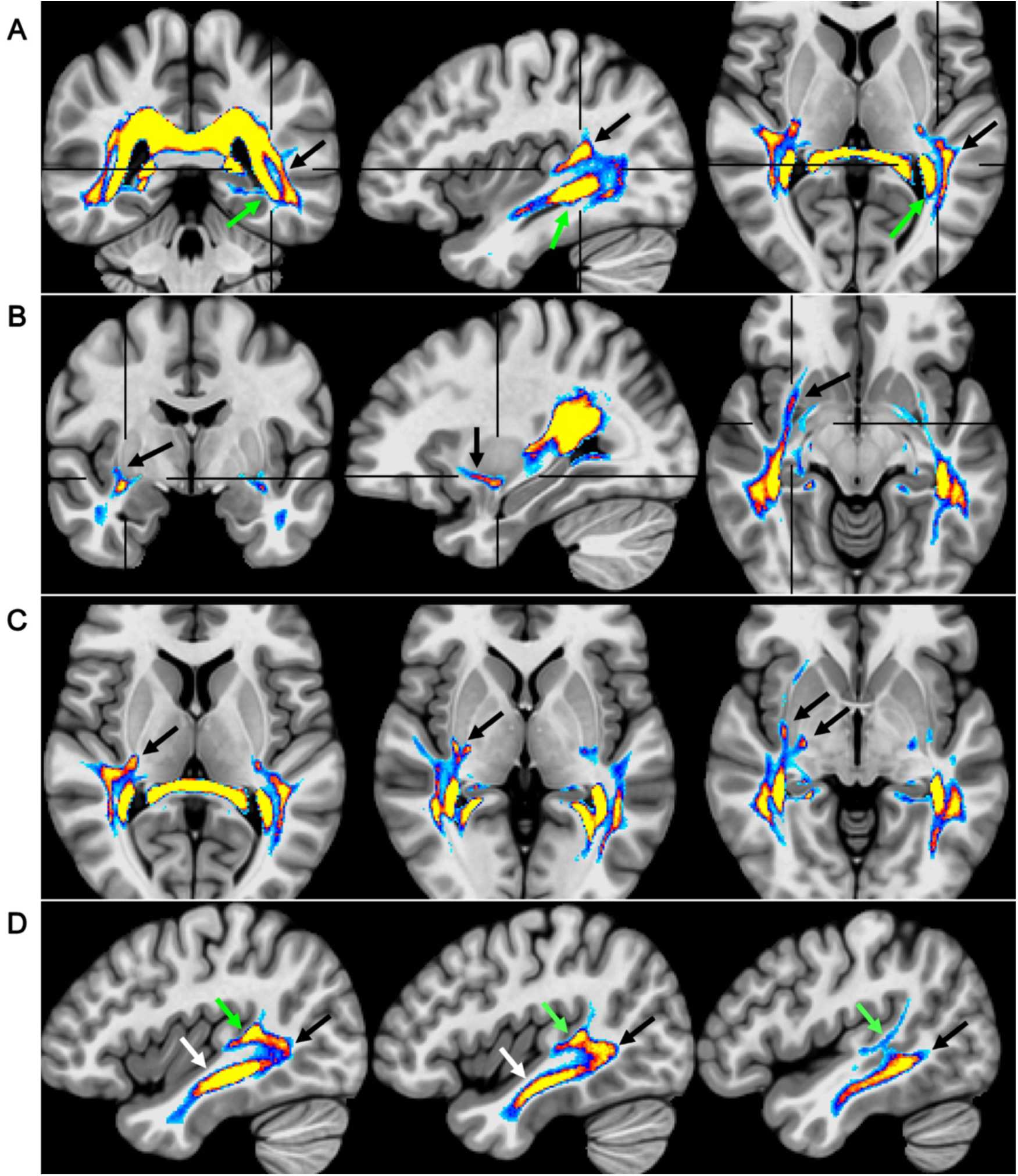
Some notable features of the transcallosal pathway. Black crosshairs show the location of orthogonal slices within each row. The left hemisphere is on the reader’s right in coronal and axial images. (A) Division of the pathway into lateral (black arrows) and medial (green arrows) divisions. (B) External capsule/claustrum fibers branching from the medial division. (C) Retrolenticular internal capsule and external capsule fibers branching from the lateral division. (D) Division of the lateral division into superior (green arrow) and inferior (black arrow) branches. White arrows in row (D) indicate the medial division, which lies just medial to the inferior branch of the lateral division.

The lateral division also arcs anteriorly from the division point above the atrium, maintaining a more superior position compared to the medial division. It courses initially through white matter deep to the supramarginal gyrus (SMG), sending a branch anteriorly into retroinsular white matter, which divides into a retrolenticular internal capsule branch and an external capsule branch that merges with the external capsule/claustrum projections from the medial division (**Figure 4c**). As it enters the white matter of the posterior superior temporal gyrus (STG), the lateral division divides further into a superior and an inferior branch (**Figure 4d**). The superior branch courses through the posterior STG just beneath the planum temporale, extending anteriorly to the base of Heschl’s gyrus and posteriorly into the SMG. The inferior branch initially courses caudally through white matter in the posterior STG (best appreciated in the sagittal plane, Figure 4d) and into the posterior middle temporal gyrus (MTG), then abruptly turns anteriorly into white matter deep to the MTG and fundus of the inferior temporal sulcus. From there it continues into deep anterior temporal white matter just lateral to the anterior extension of the medial division.

### Anterior Commissure Connections

A map of anterior commissure connections with the temporal lobe is shown in **Figure 5**. Lateral to the anterior commissure, fiber tracts traverse the Gratiolet canal in a posterolaterally oriented arc at the ventral aspect of the putamen/globus pallidus border and enter the temporal stem, as previously described (Baudo, Colombo et al. 2022; Catani and Thiebaut de Schotten 2012). Sparse fibers arising from the canal can be seen entering the ventral basal ganglia, more prominently in the right hemisphere in our sample. The tracts then spread anteriorly and posteriorly in the stem and white matter core of the temporal lobe, forming an inferolaterally oriented “sheet” extending from deep white matter lateral to the temporal horn, through the temporal stem, and superiorly into the ventral putamen and posterior subinsular white matter. Anteriorly, connections course around the lateral aspect of the amygdala and project to the temporal pole, and to the perirhinal region and anterior collateral sulcus. The posteriorly oriented tracts continue as an expanding, medially concave sheet extending from the ventromedial temporal white matter inferiorly to inferior parietal white matter superiorly. The posterior projection sends small branches to the ventrolateral thalamus, as previously described (Baudo, Colombo et al. 2022). Further posteriorly, the sheet gradually contracts in size as posterior fibers continue into the deep occipital lobe white matter lateral to the occipital horn of the ventricle.

**Figure 5.**
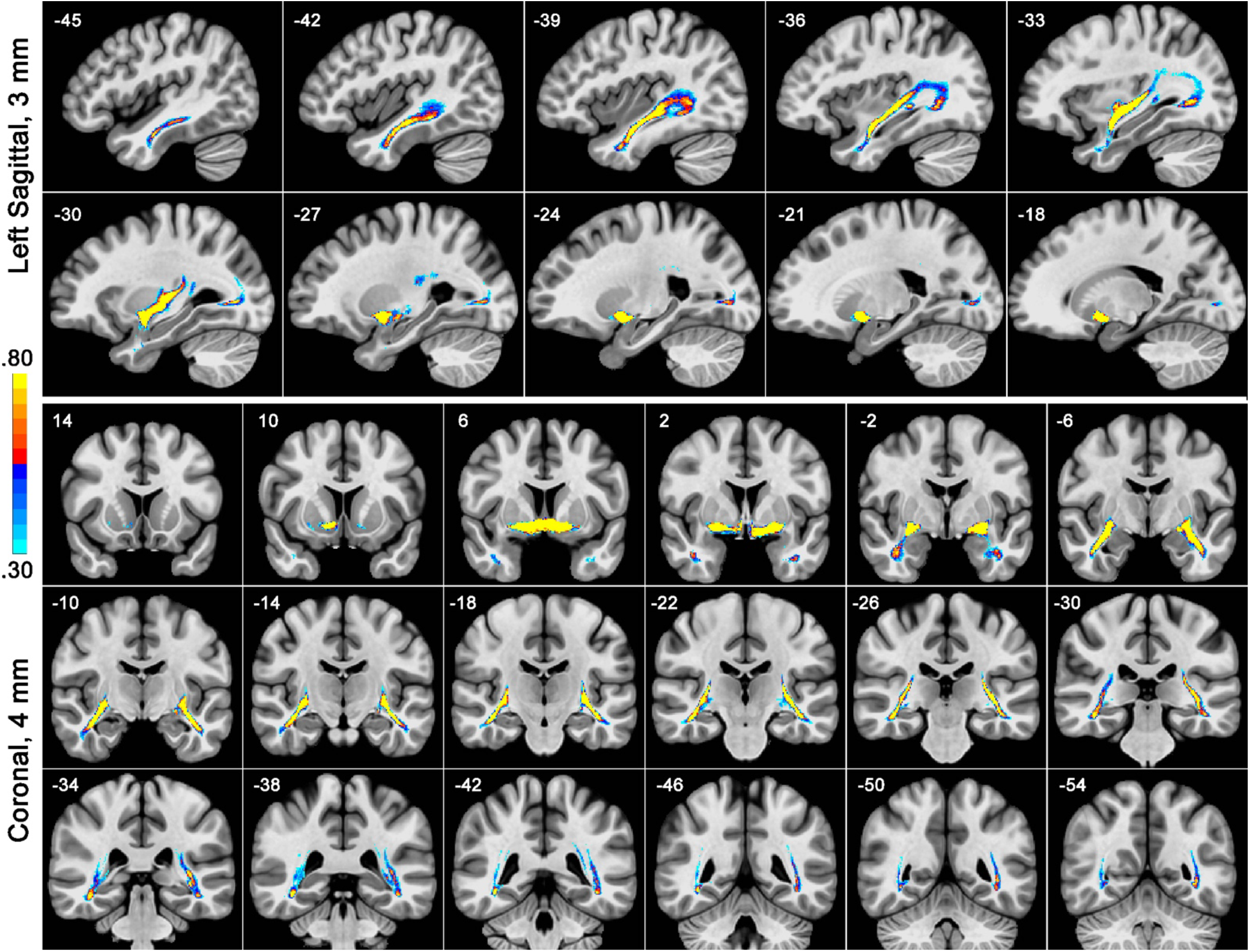
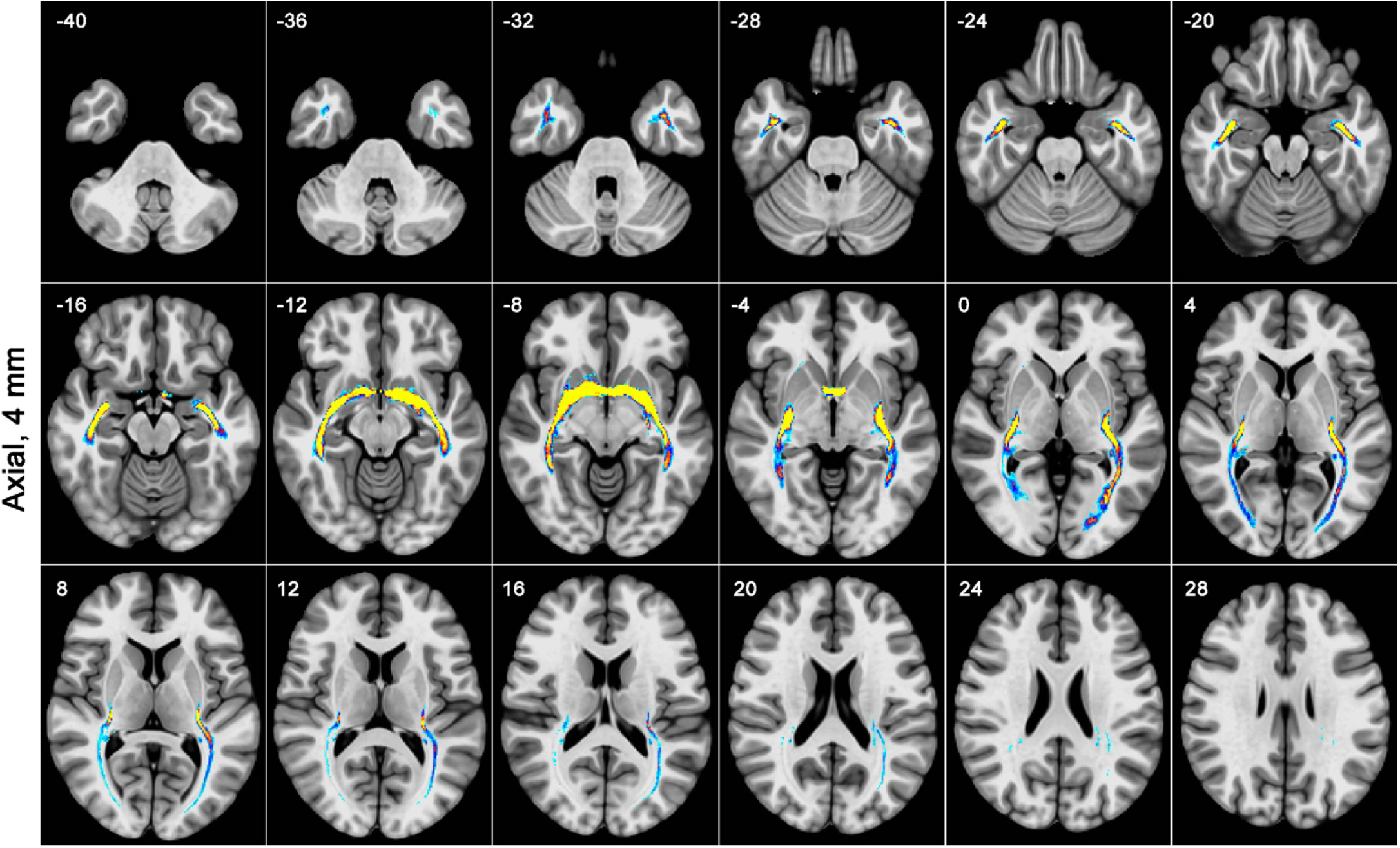
part 1. Serial sections through the averaged group anterior commissure tractography map, including left sagittal, coronal, and axial views. Formatting as in Figure 2.

### Transcallosal/Commissural Overlap

Spatial overlap between transcallosal and commissural fibers is illustrated in **Figure 6**. Major areas of overlap include much of the sheet extending from white matter lateral to the temporal horn of the ventricle, through the central temporal stem, into white matter beneath the posterior insula and inferior parietal lobe. Anteriorly, commissural fibers are located more medially in this sheet compared to the more laterally located transcallosal fibers, with overlap in the center of the sheet (**Figure 7a)**. Lateral to the tapetum, the anterior commissure tracts notably occupy white matter sandwiched between the two major divisions of the transcallosal projection (**Figure 7b**).

**Figure 6.**
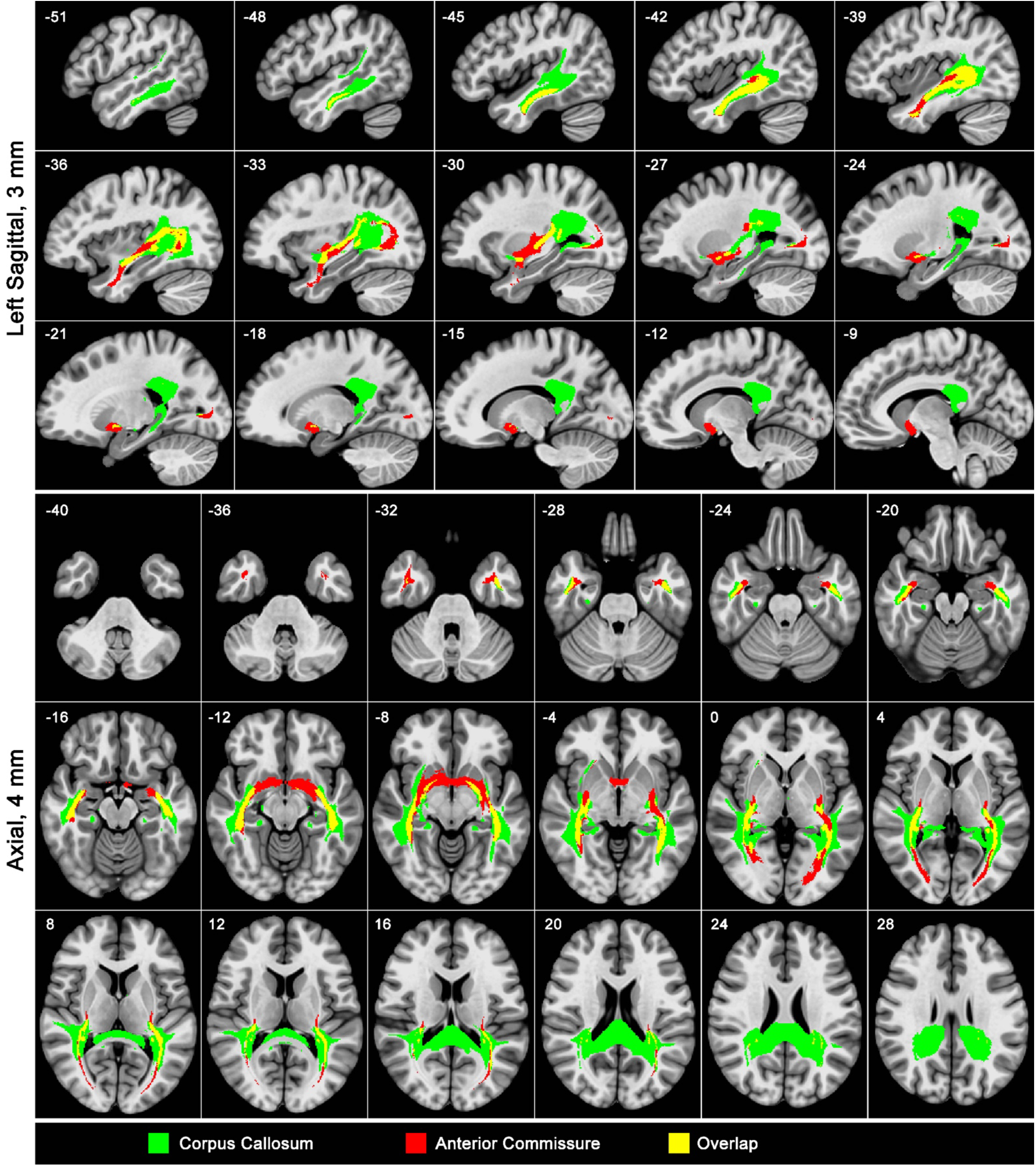
Spatial overlap (yellow) of the transcallosal (green) and anterior commissure (red) pathways.

**Figure 7.**
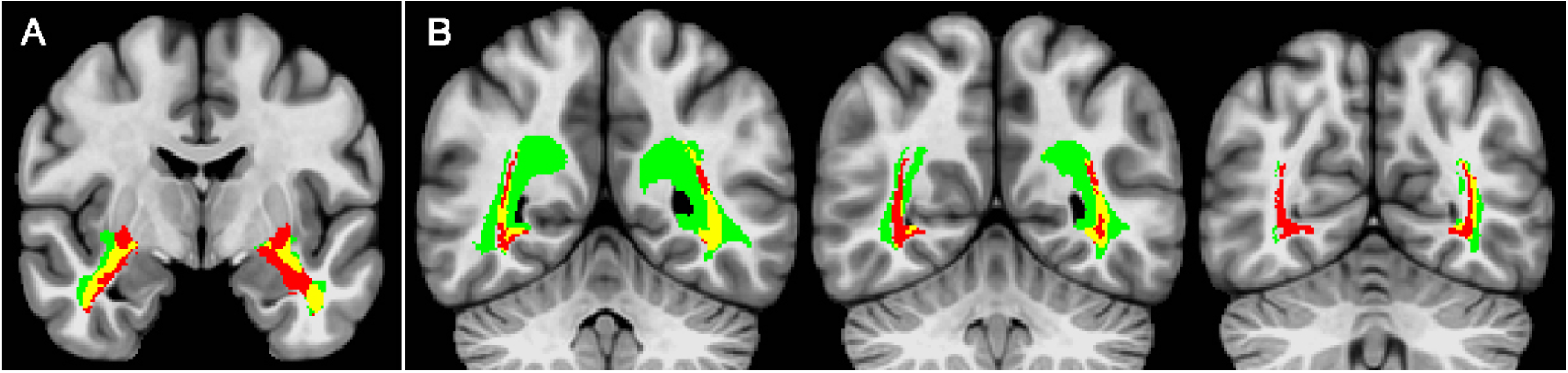
Two notable spatial relationships between the transcallosal (green) and anterior commissure (red) pathways. Overlapping areas are shown in yellow. (A) Anterior commissure fibers lie medial to transcallosal fibers in the deep temporal lobe white matter. (B) Anterior commissure fibers interdigitate between the medial and lateral divisions of the transcallosal pathway lateral to the atrium and occipital horn.

### Cortical Targets

Figure 8. shows points of intersection between individual tractography maps and individual cortical surface models, averaged on a common surface template. Cortical endpoints of the transcallosal projections appeared heaviest in the MTG and inferior temporal sulcus bilaterally (more extensive on the left and extending into the temporal pole bilaterally), the left superior temporal sulcus (STS), the posterior occipitotemporal sulcus bilaterally (more extensive on the left), and the inferior insula and posterior planum temporale bilaterally (more extensive on the left). Other areas with relatively high projection density included the anterior fusiform gyrus bilaterally, the ventromedial temporal pole bilaterally, a focal region of the right parietal STS, and the right anterior STS. High probability values in the hippocampus bilaterally likely reflect spatial proximity of the medial division (in the sagittal stratum) to dorsal-medial aspects of the hippocampus, and other labeling just lateral to the hippocampus likely represents parahippocampal projections from the DHC.

**Figure 8.**
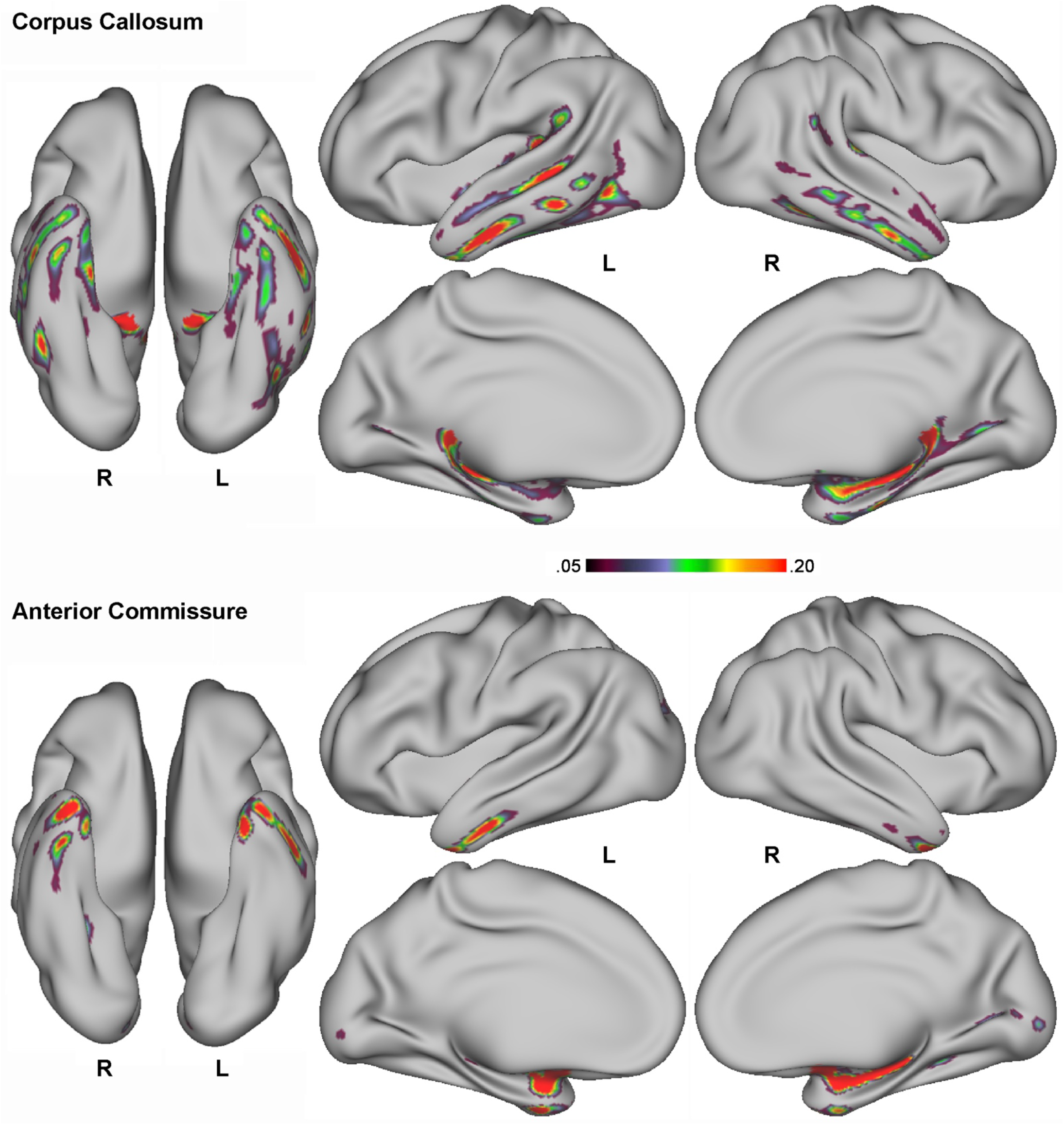
Average probability maps of cortical termination of the transcallosal and anterior commissure pathways. The corpus callosum map includes parahippocampal projections from the dorsal hippocampal commissure.

Cortical endpoints of the anterior commissure projections were heaviest in the anterior and medial temporal pole bilaterally and in left anterior fusiform gyrus. Additional projections were observed in the calcarine sulcus bilaterally, the right collateral sulcus, and the left posterior intraparietal sulcus.

## Discussion

The results provide new details about the spatial location of interhemispheric tracts connecting the left and right temporal lobes. This information has potential relevance to a broad array of processes in which the temporal lobes play a central role, including speech perception and language, semantic and episodic memory, social cognition, and emotion processes. We provide details on the location of temporal lobe projections within the splenium and find that the caudal third of the splenium contains little or no temporal lobe connections. We report a previously undocumented separation of the transcallosal pathway into distinct medial and lateral tracts and describe the course of each tract. Other details not previously described, to our knowledge, include projections from the transcallosal tract into the claustrum/external capsule, and interdigitation of transcallosal and anterior commissure projections in the deep parietal and occipital white matter. We demonstrate areas of spatial overlap between the transcallosal and anterior commissure pathways. Finally, we provide the most detailed information to date regarding the principal areas of cortical termination of interhemispheric temporal lobe connections.

The distinct location of the medial and lateral transcallosal tracts suggests that they have different functional roles. The medial tract, for example, is unlikely to play a role in speech perception, given that it probably does not project to superior or lateral temporal regions where these processes are supported. Its location in the medial white matter instead suggests a role in episodic memory, semantic processing, or emotional cognition. It shows extensive spatial overlap with the posterior projection from the anterior commissure, the two tracts together forming a larger anterior-posterior medial temporal network.

The lateral transcallosal tract is likely to be involved in speech perception and language processes given its projections to posterior superior temporal (especially planum temporale), middle temporal, and superior temporal sulcus regions. A lesion that simultaneously damages the left-lateralized phoneme perception system and interrupts transcallosal input to left hemisphere language networks would effectively deprive the language system of auditory verbal input, thereby causing a relatively isolated speech comprehension deficit (with intact reading comprehension) or contributing to comprehension impairments in cases of Wernicke aphasia. Previous authors have proposed this “interhemispheric disconnection” mechanism to explain cases of “pure word deafness” from unilateral left hemisphere damage, analogous to the high-level visual disconnection that underlies some instances of “pure alexia”, though the relevant lesion location is not clear (Takahashi, Kawamura et al. 1992, Poeppel 2001, Maffei, Capasso et al. 2017). Our data suggest that a focal lesion damaging the left superior temporal sulcus, the main area implicated in phoneme perception (Liebenthal, Binder et al. 2005, Hickok and Poeppel 2007, Binder 2015), or the mid-MTG, together with damage to the lateral transcallosal tract transmitting phonemic information from the right hemisphere, could result in such a disconnection. Likely sites of damage to the lateral transcallosal tract include white matter deep to the planum temporale and SMG, affecting the superior branch of this tract, and white matter in the middle to posterior MTG, affecting the inferior branch. Some current language models suggest that the planum temporale and SMG play a larger role in speech production than in speech perception (Hickok and Poeppel 2007, Binder 2015), suggesting that interhemispheric projections to this region could be less critical for phoneme perception than those projecting more ventrally to the STS and MTG.

The diffusion tractography methods used here are limited in their ability to track projections into the cortex, and the cortical projection maps we provide here should be considered alongside that caveat. They likely detect the few fibers that are close enough to the cortex to show overlap with the cortical ribbon within the relatively coarse spatial grid of diffusion tensor imaging. We nevertheless believe they provide useful new information on likely termination patterns of these interhemispheric pathways.

## Statements and Declarations

### Author Contributions

Jeffrey R. Binder conceived and designed the study. Jedediah Mathis, Sidney Schoenrock, and Volkan Arpinar prepared the materials and carried out the data collection. Mónica Giraldo-Chica, Jedediah Mathis, Jia-Qing Tong, and Joseph Heffernan carried out the data analysis. The first draft of the manuscript was written by Jeffrey R. Binder, and Mónica Giraldo-Chica and L. Tugan Muftuler commented on previous versions of the manuscript. All authors read and approved the final manuscript.

### Data Availability

The datasets acquired and analyzed for this study are available from the corresponding author on request.

### Ethics Approval

The project was approved by the MCW Institutional Review Board (Protocol 00024807, approval date 8/31/2015) and was performed in accordance with the Declaration of Helsinki.

### Consent to Participate

All participants provided written informed consent for the study.

## Acknowledgments

Supported by grants from NINDS (U01 NS093650), Advancing a Healthier Wisconsin Foundation (Project# 5520462), and GE Healthcare. The authors thank Sarah Smith for help with participant recruitment and scheduling.

